# Purging due to self-fertilization does not prevent accumulation of expansion load

**DOI:** 10.1101/2022.12.19.521096

**Authors:** Leo Zeitler, Christian Parisod, Kimberly J. Gilbert

## Abstract

Species range expansions are a common demographic history presenting populations with multiple evolutionary challenges. It is not yet fully understood if self-fertilization, which is often observed at species range edges, may create an evolutionary advantage against these challenges. Selfing provides reproductive reassurance to counter Allee effects and selfing may purge accumulated mutational burden due to founder events (expansion load) by further increasing homozygosity. We study how selfing impacts the accumulation of genetic load during range expansion via purging and/or speed of colonization. Using simulations, we disentangle inbreeding effects due to demography versus due to selfing and find that selfers expand faster, but still accumulate load, regardless of mating system. The severity of variants contributing to this load, however, differs across mating system: higher selfing rates purge large-effect recessive variants leaving a burden of smaller-effect alleles. We compare these predictions to the mixed-mating plant *Arabis alpina*, using whole-genome sequences from refugial outcrossing populations versus expanded selfing populations. Empirical results indicate accumulation of expansion load along with evidence of purging in selfing populations, concordant with our simulations, and suggesting that while purging is a benefit of selfing evolving during range expansions, it is not sufficient to prevent load accumulation due to range expansion.

**Author Summary:** The geographic space that species occupy, i.e. the species range, is known to fluctuate over time due to changing environmental conditions. Since the most recent glaciation, many species have recolonized available habitat as the ice sheets melted, expanding their range. When populations at species range margins expand into newly available space, they suffer from an accumulation of deleterious alleles due to repeated founder effects. We study whether self-fertilization, which is considered an evolutionary deadend, can be favored under these expanding edge conditions. Selfing has two important effects: allowing for faster expansion due to reproductive assurance and purging recessive deleterious alleles by exposing them to selection as homozygotes. We use simulations to identify the impact of selfing on expanded populations and then compare these results to an empirical dataset to assess whether our predictions are met. We use the mixed-mating plant alpine rock-cress (*Arabis alpina*) since it has both expanded since the last glaciation and undergone a mating shift to selfing. We find that selfing does not prevent the accumulation of deleterious load, however purging does still act to remove the most severe variants, indicating that selfing provides this benefit during range expansions.

## Introduction

Many species across the globe have expanded or shifted their species ranges in response to changing climates (Davis and Shaw 2001; Parmesan 2006), creating unique evolutionary and demographic conditions. The repeated bottlenecks and founder events that characterize populations at expanding range edges increase the strength of genetic drift, reducing genetic variation, and reduce the efficacy of selection (Caballero 1994; Wright 1931). The study of species range expansions thus provides interesting insight into evolutionary questions of if and how populations manage to adapt and continue spreading despite reduced genetic diversity as well as other difficulties faced at range fronts. Individuals colonizing previously unoccupied environments face limited mate or pollinator availability, resulting in slower expansion as a result of these Allee effects (Moeller *et al*. 2012; Dennis 1989; Courchamp *et al*. 1999; Stephens and Sutherland 1999; Hallatschek and Nelson 2008). Gene surfing, whereby variants increase in frequency at expanding edges due to serial founder events (Burton and Travis 2008; Hallatschek and Nelson 2010; Edmonds *et al*. 2004; Klopfstein *et al*. 2006), can also affect selected variants due to the reduced efficiency of selection. The surfing of deleterious variants at an expanding edge is thus possible, and this process has been shown to lead to the accumulation of deleterious variants in expanded populations, causing what is termed expansion load (Peischl *et al*. 2013, 2015; Peischl and Excoffier 2015). Expansion load has the potential to temporarily halt population growth or cause local extinction at the boundaries of the species range (Peischl *et al*. 2013, 2015; Gilbert *et al*. 2017, 2018). Evidence of elevated load due to expansion is well-documented empirically, including in humans (Henn *et al*. 2015, 2016; Peischl *et al*. 2018), plants (González-Martínez *et al*. 2017; Willi *et al*. 2018), and experimental bacterial populations (Hallatschek and Nelson 2010; Bosshard *et al*. 2017). Whether some organisms are able to overcome the burden of expansion load through adaptive measures during expansion is, however, unclear.

Numerous plant species are known to have expanded their ranges after the last glacial maximum, and it is widely observed that many of these species exhibit a transition to self-fertilization (‘selfing’) at range edges (Barrett 2002). Selfing can reduce fitness through the expression of inbreeding depression (Charlesworth and Charlesworth 1987; Reusch 2001; Barrett 2013), and is often considered to be an evolutionary dead end, as it can lead to mutational meltdown (Lynch *et al*. 1995; Goodwillie *et al*. 2005; Igic and Busch 2013). Yet the observation of enrichment for selfing at species range edges suggests that there is some advantage gained with this mating system under the demographic and evolutionary conditions that range expansions impose. Simulations have likewise predicted that selfing should evolve at species range edges during expansion (Encinas-Viso *et al*. 2020).

Selfing has three major evolutionary impacts on populations: reducing effective population size, reducing the effective recombination rate, and removing or reducing Allee effects. Selfing can remove Allee effects by assuring reproductive success in low density populations (Baker 1967; Lloyd and Schoen 1992), and therefore lead to faster expansion speeds, already evidenced by some studies of organisms with uniparental reproduction (Pannell and Barrett 1998; Eriksson and Rafajlovic’
s 2021). This is a clear and expected advantage during species range expansion, as colonization can be faster. However, the reduction in *N*_*e*_ along with reduced effective recombination rate due to selfing should be disadvantageous and exacerbated when compounded with the already reduced genetic diversity due to founder effects during range expansion. Some evidence suggests that range expansions may instead facilitate a transition to selfing by depleting the genetic load at the edge and reducing inbreeding depression (Pujol *et al*. 2009). Because selfing greatly increases homozygosity, it creates the potential for exposing recessive deleterious alleles to selection and thus purging them from the population (Ohta and Cockerham 1974; Charlesworth 1992; Glémin 2007; Pujol *et al*. 2009). Purging therefore has the potential to counteract the accumulation of expansion load. Though, Glémin (2007) concluded that purging might only be a short term effect of selfing, and over long evolutionary time scales fixation would be the dominant effect in selfers. Evidence suggests that in small populations purging by self-fertilization is less feasible, and small-effect deleterious variants can still contribute to an increase in genetic load (Wang *et al*. 1999). The distribution of selection coefficients compared to population size is thus a major factor for the evolution of mating systems (Bataillon and Kirkpatrick 2000; Glémin 2003). As a consequence, one important change in the population genetic signature of a transition from outcrossing to selfing is a shift in the observed distribution of fitness effects (DFE) for variants segregating in the population to reflect less efficient selection against weakly deleterious, additive variants and purging of strongly deleterious, recessive variants (Barrett *et al*. 2014; Laenen *et al*. 2018; Arunkumar *et al*. 2015). Whether this prediction holds when a species range expansion occurs concurrently with a mating system shift has, to our knowledge, not been fully explored.

In combination with simulations of a range expansion and mating system shift, we additionally investigate an empirical system matched to this demographic and evolutionary history. The perennial arcticalpine plant *Arabis alpina* L., is a species known to have been subject to range expansions and contractions in response to the repeated quaternary climate oscillations (Ansell *et al*. 2008). The Italian peninsula is a known refugium for outcrossing populations during the last glacial maximum (Ansell *et al*. 2008), after which the species recolonized alpine habitats across Europe and concurrently with this expansion evolved a mating system of predominant self-fertilization (Tedder *et al*. 2015). In Italy, populations of *A. alpina* are predominantly outcrossing with high genetic diversity (Ansell *et al*. 2008; Laenen *et al*. 2018), while in the French and Swiss Alps, populations are more homozygous with lower nucleotide diversity, and mainly selfing (Ansell *et al*. 2008; Buehler *et al*. 2012). Given its demographic history of range expansion and the variation in outcrossing rates, we study the combined effect of a range expansion and an evolutionary transition in mating system from natural populations across Italy and the western Alps.

In this study, we test the hypothesis that selfing may be evolutionary favored during range expansions due to the ability to purge otherwise accumulating deleterious expansion load. We investigate how selfing at the range edge affects genetic diversity, mutation load, the observed DFE, and the speed of colonization during a range expansion. Using individual-based simulations, we model a range expansion where different degrees of self-fertilization are introduced mid-expansion and compare to a null model of obligately outcrossing expanding populations. We focus in particular on the dynamics of load accumulation with evolved selfing rates and include lethal, mildly deleterious, and beneficial mutations with different dominance coefficients. By characterizing the distribution of selection coefficients of expanded populations, we highlight differences in the efficacy of selection across mutation severity. We also compare the buildup of genetic load across selfing rate scenarios and estimate load under different assumptions of dominance. We then test if our predictions from simulations are equivalently detectable in empirical data from natural populations of *A. alpina* in the Italian-Alpine expansion zone. We find that rapid purging due to self-fertilization is common for highly deleterious recessive variants but does not prevent the accumulation of genetic load in expanded populations. Our results mark a vital step towards deciphering the interplay of complex population demography and mating system on the fate of genetic diversity and mutation load, helping us to better understand how populations react to changing environmental conditions.

## Results

### Selfing leads to faster expansion

Using individual-based, forward-time simulations in SLiM v3.7.1 (Haller and Messer 2019), we modelled a range expansion across a one-dimensional linear landscape. We simulated obligate outcrossing from the first deme (‘core’) followed by a shift to self-fertilization in the 25th deme of the landscape (out of 50 total demes), with selfing rates *σ* of 0.5, 0.95, or 1, or for a null comparison, simulations with continued obligate outcrossing across the entire landscape during expansion and colonization (Figure 1A).

**Figure 1:**
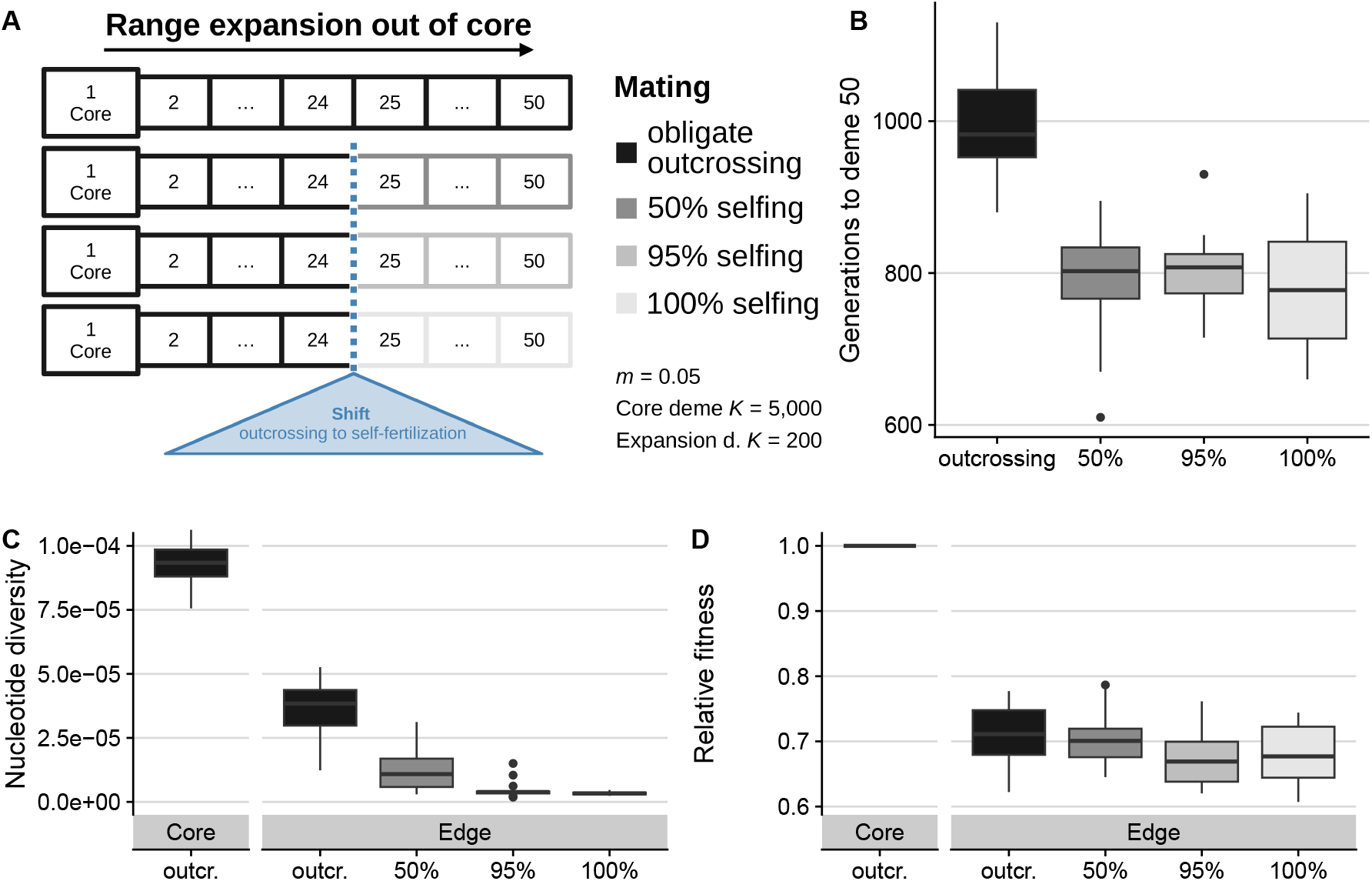
Simulation schematic for 1-D landscapes with stepping-stone migration (A). A shift in the rate of self-fertilization occurs in the center of the landscape (blue triangle). The number of generations needed to cross the landscape from the core to deme 50 (B). Expansion time was lower for all selfing rates but higher for obligate outcrossing. Mean nucleotide diversity (C) was reduced outside of the core, with a greater reduction for higher selfing: 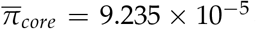, 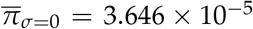, 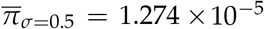, 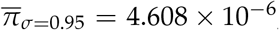, 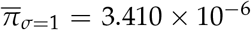. Relative fitness (D) decreased from core to edge with similar values across outcrossers and selfers: 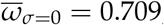, 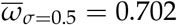, 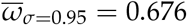, 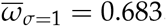, reflecting a relative loss of fitness as compared to the core of 29.1%, 29.8%, 32.4%, and 31.7% respectively for *σ* = 0, 0.5, 0.95, and 1.

One expected benefit of selfing during range expansion is increased expansion speed, which we observed in our simulation results. We compared the number of generations required to cross the landscape among selfing rates and found that mixed mating and obligate selfing populations had a faster expansion speed compared to obligate outcrossing populations (see Figure 1B). The differences in generation time among different selfing rates was minor (mean generation times for *σ* = 0.5, 0.95, and 1, respectively: 792 (*SD* = 78.5), 800 (*SD* = 51.4), 777 (*SD* = 70.7)) compared to the notable difference in expansion time of obligate outcrossers (990 generations (*SD* = 69.8)).

### Range expansion increases genetic load and decreases diversity in simulations

To test if and how selfing modifies the outcomes of a range expansion in our simulations, we examined genetic diversity in expanded populations and across selfing rates. Both outcrossing and selfing edge populations showed large reductions in diversity due to the expansion. Outcrossers retained the highest nucleotide diversity for neutral sites (*π*_*edge,σ*=0_ = 3.646 × 10^−5^), while with increasing self-fertilization rates, populations showed further reductions in nucleotide diversity (*π*_*edge,σ*=1_ = 3.410×10^−6^, Figure 1C). Core populations which never experienced expansion and always outcrossed had the highest nucleotide diversity (*π*_*core*_ = 9.235 × 10^−5^).

Genetic load is predicted to be higher in expanded populations, so we next examined how selfing modulates this outcome of a range expansion. With simulations we could accurately distinguish inbreeding effects due to mating of related individuals in small populations at the range edge versus inbreeding effects resulting from uni-parental inheritance, *i*.*e*. selfing, by contrasting obligate outcrossing scenarios to those with various rates of selfing. Fitness of every individual is also known, as this is defined in SLiM as the target number of offspring to be generated by an individual and is calculated multiplicatively across the effects of all derived mutations (see Methods for a full description). We calculated mean fitness of all individuals within a deme per replicate 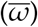 and compared these values between core and edge populations after the simulated expansion was complete. In all cases, the range expansion reduced fitness at the edge due to expansion load (Figure 1D). In the obligate outcrossing case (*σ* = 0), we observed a reduction of fitness from core to edge of 29.1%. Interestingly, selfers showed negligible differences in load accumulation relative to outcrossers, with at most a mean reduction in fitness of 31.7% for obligate selfers. We observed an increase in the proportion of loci fixed for deleterious alleles in all expanded populations, with these proportions increasing for higher selfing rates (Figure S1A). Similarly, we found that mean counts of deleterious loci increased from core to edge as well as from lower to higher rates of selfing (Figure S1B), whereas counts of deleterious alleles showed less to no clear pattern from core to edge and among selfing rates (Figure S1C).

To understand why and how self-fertilization seemingly had no impact on removing genetic load during a range expansion, we examined demes over time and space to disentangle the effects of inbreeding due to demography versus inbreeding due to increased self-fertilization (Figure 2). Outcrossing deme 24 exhibited a mean observed heterozygosity level of 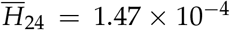when it was first colonized during the expansion, i.e., when it was the edge of the species range. Beyond deme 24 the mating system shifts to selfing and we observed a continual loss of heterozygosity at the expanding front. When the expansion front reached deme 35, 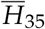 for 95% selfers rapidly decreased to 7.05 ×10^−7^. For outcrossers, however, heterozygosity exhibited a more gradual rate of reduction across the course of expansion, reach-ing 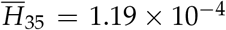by the time deme 35 was colonized (Figure 2A, S2). We examined how diversity recovered in deme 35 over time since its colonization until the end of the simulation and found that outcrossers recovered to higher levels (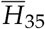 at the end of the simulation 2.27 × 10^−4^) than selfers (for 95% selfing 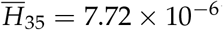.

**Figure 2:**
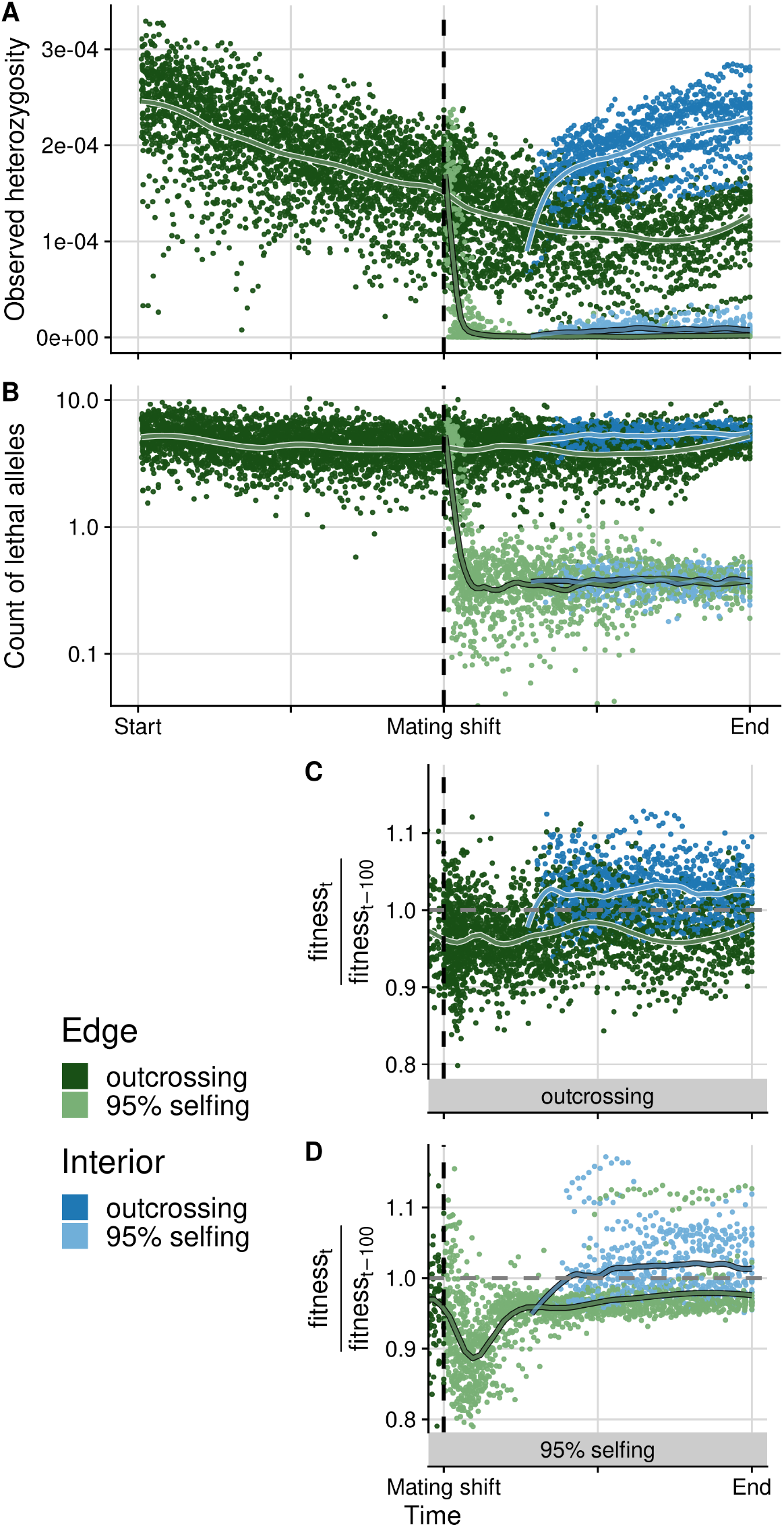
Observed heterozygosity (A), count of lethal alleles (B), and the rate of fitness change (C-D) are shown through time for the expanding range edge (green) as compared to change over time within one interior deme, stationary on the landscape (blue, deme 35). For selfing rates 50% and 100%, see Figures S2 and S3. Panels (C) and (D) show the rate of fitness change as measured over 100-generation intervals, separately for outcrossers and selfers. A value of 1 indicates no change in fitness over 100 generations while values above 1 indicate increasing fitness and values below 1 indicate fitness loss. The vertical dashed line indicates the point in time where the mating system shifts to selfing. This shift occurs at deme 25 on the landscape, and since there is variation across simulation replicates in the generation time taken to expand to deme 25, we plotted all values relative to this time point for each replicate shown (*n* = 20 replicates per selfing rate scenario). Each point is the value from a single simulation replicate and lines are loess (span=0.2) fitted curves across all replicates.

### Fitness loss despite genetic purging

We counted the number of lethal alleles per individual and observed a reduction in the count of lethals that corresponds with the reduction in heterozygosity (Figure 2A-B). Lethal alleles were only reduced when the shift to selfing occurred, and we observed the same pattern at every simulated selfing rate. Obligate outcrossers did not exhibit a reduction in lethal alleles, showing no evidence of purging. The largest reduction in lethal alleles occurred for the shift to the highest selfing rate (*σ* = 1) with a 91.69% drop in lethal alleles, while our lowest simulated selfing rate (*σ* = 0.5) still showed a strong effect of purging lethal alleles with an 83.74% reduction (Figure S3).

The rate of change of fitness in a given deme, measured at a focal generation (*t*) compared to 100 generations prior (*t*− 100), showed a consistent loss of fitness over time due to range expansion as well as some fitness recovery in populations behind the expanding front (Figure 2C, D). Edge demes which recently underwent the shift to selfing exhibited a drastic reduction in fitness relative to equivalent outcrossers. This high rate of fitness loss exhibited by selfers is temporary and only lasts for between 45-235 generations, after which the rate of fitness loss recovers to the same rate as that observed in expanding obligate outcrossers: still below one on average and accumulating expansion load.

We also investigated the impact the range expansion and mating shift had on the realized distribution of selection coefficients. Overall, we found the greatest proportion of deleterious mutations in the weak to intermediate bin of selection coefficients (−0.0001 ≤ *s* < −0.001), with just below 60% of all sites falling into this class (Figure 3). The next most deleterious bin (− 0.001 ≤ *s* < − 0.01) contained about 30% of sites, while about 10% of sites are in the weakest selection coefficient bin. Lethal alleles made up a small proportion of segregating sites, as expected given the small proportion defined in the simulation parameters. Within these small numbers of severely deleterious variants, there was a consistent trend for a reduction of lethals from core to edge of nearly 50% for outcrossers and significantly further reduction for all rates of selfing, increasing from a nearly 75% reduction from core to edge for 50% selfers to more than 75% for 100% selfers (Figure 3 inset). The pattern of reduced proportions of deleterious sites as selfing rate increases holds in both the lethal category as well as the second-most deleterious allele class. We consistently observed the reverse pattern in the remaining weaker effect bins, with proportions of weakly deleterious sites slightly increasing at range edges and more so with higher selfing rates. This observation is consistent with more efficient removal of highly deleterious alleles and mutation accumulation at sites with smaller absolute selection coefficients.

**Figure 3:**
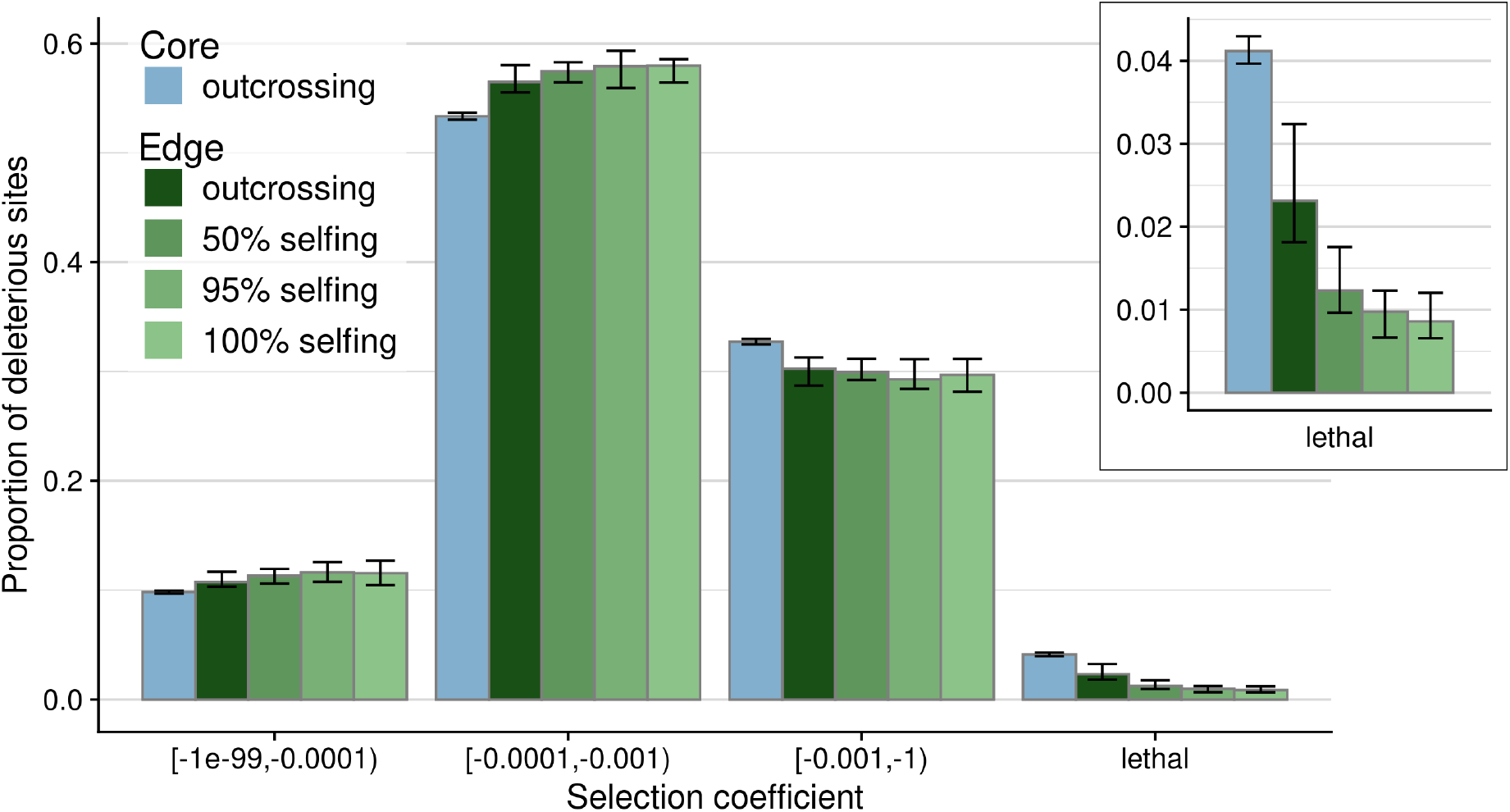
The observed distribution of selection coefficients from simulations at the end of expansion. The core deme (blue) is compared to edge demes (green) for obligate outcrossing (darkest color) versus higher selfing rates (lighter colors). Error bars indicate 0.05 and 0.95-quantiles across the 20 simulation replicates. The inset panel emphasizes the degree to which the proportion of sites in the lethal category changes over mating system scenario from core to edge.

### Reduced genetic diversity and elevated load in expanded selfing *A. alpina* populations

To test if our observations for genetic diversity and load accumulation from simulations are matched in natural populations, we used the mixed mating plant *Arabis alpina*, which underwent a range expansion concurrently with a shift to higher self-fertilization rates from Italy (outcrossing, Tedder *et al*. (2015)) into the Alps (selfing, see Figure 4A). Using 191 newly sampled and sequenced short-read genomes from Italy and France combined with publicly available data from Switzerland and across Europe (Laenen *et al*. 2018; Rogivue *et al*. 2019), we examined differences across the species range in 527 individuals at high resolution. Population structure results showed expected clustering by regions (Figure S4), matching the geography of sampled populations from Abruzzo in southern Italy, the Apuan Alps in northern Italy, the French Alps, and the Swiss Alps (Figure 4A). Previously sampled individuals from Italy, France, and Switzerland that we combined with our newly sampled individuals also consistently clustered within the same geographic regions. Samples of fewer individuals from more widely across Europe showed reasonable structuring among Greek, Spanish, and Scandinavian populations. Inference of the demographic history of the Italian and alpine populations populations showed a history of population bottlenecks and recovery consistent with the northward expansion following the last glacial maximum and the shift to selfing in France and Switzerland (see Figure S5).

**Figure 4:**
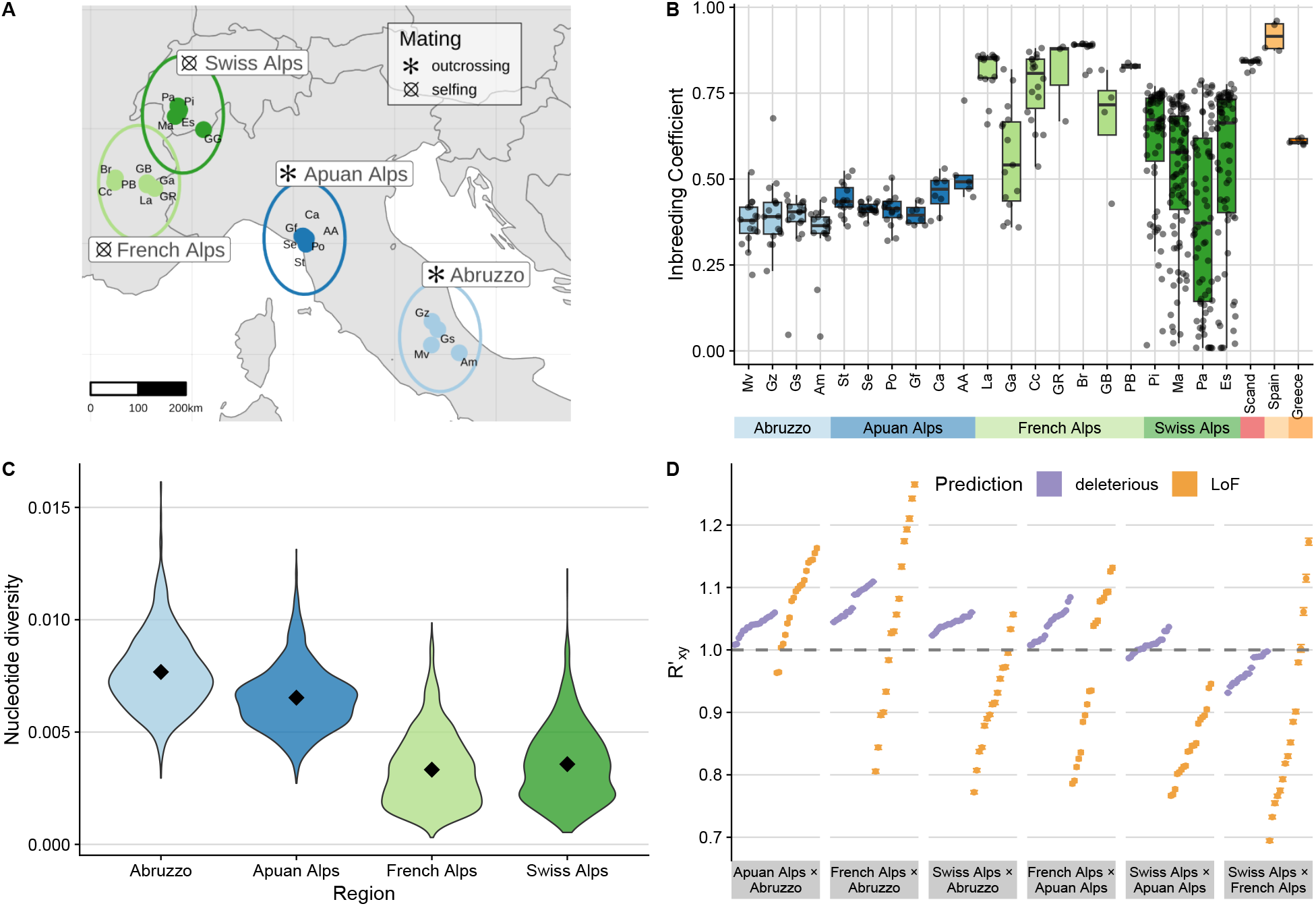
Sampling sites of *A. alpina* in the Italian-Alpine expansion zone (with mating types as published in Buehler *et al*. 2012; Tedder *et al*. 2015) (A). Inbreeding coefficients for individuals across sampled populations, including Spain (selfing), Scandinavia (selfing), and Greece (outcrossing, Laenen *et al*. 2018) (B). The distribution of nucleotide diversity estimated for Italian and alpine populations (C), with diamonds indicating group means. 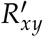 values for deleterious (purple) and LoF (orange) loci (D). 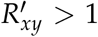 indicates an accumulation of derived alleles at deleterious or LoF sites relative to neutral ones, while 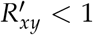 indicates the opposite.

To reveal how the expansion and self-fertilization impacts key diversity parameters, we calculated individual inbreeding coefficients (*F*) and nucleotide diversity (*π*). We found the highest inbreeding coefficients in mixed mating and highly selfing populations outside of Italy, and reduced inbreeding in the Apuan Alps and Abruzzo (means for Swiss, French, Apuan Alps and Abruzzo, respectively: 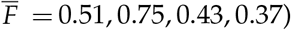. The highest overall inbreeding coefficients were estimated in populations from Spain 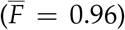 and France (Br, 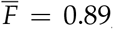, Figure 4B). Swiss populations had the greatest standard deviation(Pa, *SD*(*F*) = 0.26), and Italian populations had the lowest mean value (Am, 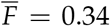). For nucleotidediversity, we found high values in the Abruzzo region of Italy (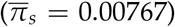). Genetic diversity reduced when moving north to the French Alps (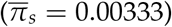) and Switzerland (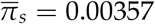, Figure 4C).

We calculated 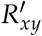to assess the accumulation of derived deleterious alleles, using 270,889 SNPs annotated as deleterious and 2380 as loss of function (LoF) variants, classified by SNPeff (Cingolani *et al*. 2012). 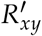 is a pairwise statistic that compares the count of derived alleles found in one population relative to another, and avoids reference bias introduced by branch shortening (Do *et al*. 2015). 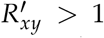 indicates that population X has more derived alleles of a given class than population Y relative to the neutral expectation, while 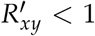 would indicate fewer derived alleles in population X. For deleterious sites we found that alpine populations had more derived alleles compared to Italian populations (Figure 4D), indicating an increase in genetic load from south to north. Within the Alps, all Swiss populations had reduced derived allele frequencies compared to France, while relative to the Apuan Alps in northern Italy, only few Swiss populations exhibited reduced derived allele counts, potentially suggesting that the shift to selfing has begun to alleviate the accumulation of expansion load. For LoF loci, signals of both purging and accumulation were detectable. Some pairwise population comparisons showed an increase in number of LoF alleles from south to north (e.g., nearly all Apuan × Abruzzo comparisons), while oth-ers showed mixed results, depending on the focal populations (e.g., French × Apuan, Swiss × French). All population comparisons of Swiss Alps × Apuan Alps showed reduced LoF allele counts.

We next used the SNPs with variants annotated as putatively deleterious to examine the accumulation of genetic load in our expanded populations. We assessed the predictability of genetic load estimation under two different assumptions of dominance: first we calculated an additive model which counted individual deleterious alleles and compared to a second recessive model which counted individual homozygous deleterious loci. In our simulations, the correlation between per-population mean fitness and load prediction was stronger for the recessive model (*R*^2^ = 0.82, *P* < 0.001) than the additive model (*R*^2^ = 0.10, *P* < 0.001, Figure S6), supporting the appropriateness of the recessive load model even in the presence of selfing. The additive model also predicted load more poorly in a supplemental set of simulations using only fully additive mutations (see Figure S7). Estimating load from these models in the empirical dataset indicated higher load in expanded, selfing populations from France and Switzerland compared to core Italian populations, using the recessive model (Figure 5A). The additive model found slightly reduced load for alpine populations compared to Italian ones (Figure 5B).

**Figure 5:**
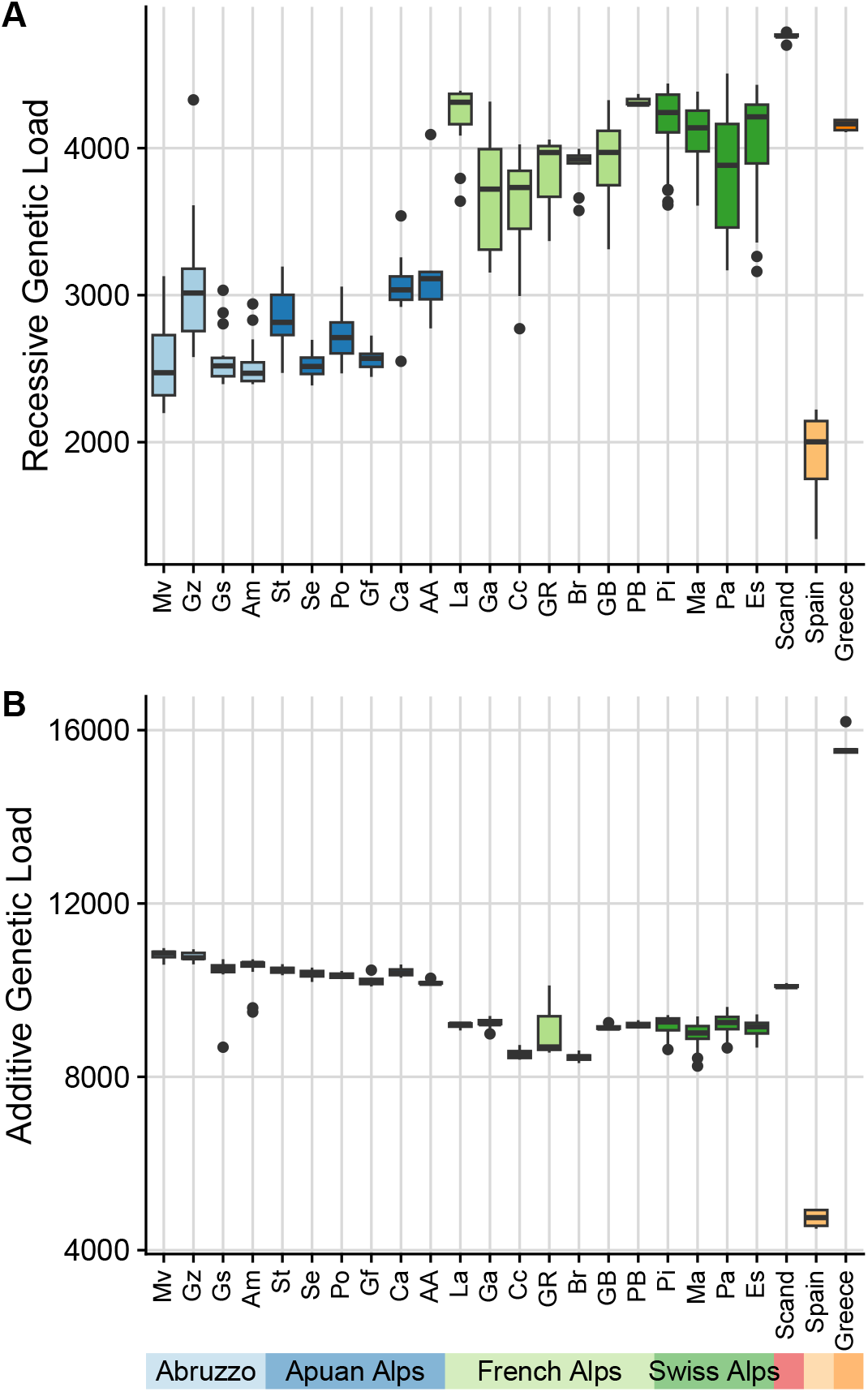
Genetic load in *A. alpina* populations as inferred from counts of deleterious loci using a recessive model (A) versus counts of deleterious alleles using an additive model (B). Loci are classified as putatively deleterious by SNPeff (see Methods).

To further understand the mutational burden within our *A. alpina* populations, we estimated the distribution of fitness effects of new mutations (DFE) using fitdadi (Kim *et al*. 2017), which corrects for demographic history by first fitting a best demographic model to the data (Figure S5). fitdadi surprisingly estimated a similar DFE across all of our sampled populations (Figure S8), with a large proportion of strongly deleterious sites at or above 60%, around 20% of sites in the weakest selection class, and approximately 5% in each of the two intermediate selection classes. The proportions varied only marginally across core Italian populations as well as across expanded French and Swiss populations. We additionally examined the fixation of deleterious alleles across our populations, within classes of neutral, deleterious, or LoF sites (Figure S9). Fixation of all sites increased from Italy in the south to France in the north, but then decreased from France to Switzerland, reminiscent of our 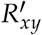 results suggesting more purging in Swiss populations.

## Discussion

In this study, we investigated the impact of selfing on the accumulation of genetic load during a species range expansion. We used simulations to disentangle the reduction in effective population size at expanding fronts due to self-fertilization versus serial founder events. We then compared our expectations for the impact of selfing to empirical data from natural populations having undergone both a range expansion and a mating system shift. Because selfing reduces the effective recombination rate within populations as well as genetic diversity, it is expected to be generally maladaptive for evolution and adaptation. However, conditions at the expanding edge of a species range may particularly favor the evolution of selfing mating systems. And the compounded effects of reduced diversity due to selfing at range edges may even provide an additional benefit of purging homozygous recessive deleterious mutations.

One clear advantage that we confirm is that selfing provides reproductive assurance (Igic and Busch 2013) and leads to faster spread over geographic space. Despite similar losses in fitness from core to range edge for both outcrossers and all selfing rate scenarios, selfers still colonized the landscape faster than outcrossers. This result adds to the general prediction of Baker’s Law, that selfing may be advantageous in mate-limited environments (Baker 1967). Since we do not have equivalently expanded outcrossing populations in our *A. alpina* dataset nor a generation time measurable in contemporary time, we cannot empirically investigate this question. To fully understand the benefits of reproductive assurance from selfing, it may be fruitful for future empirical studies to focus on organisms with well-documented expansion times and mating system shifts, or potentially take advantage of laboratory experiments with expansions of outcrossers versus selfers under controlled conditions, for example using mixed mating species of *Caenorhabditis*.

Our main interest in comparing simulation and empirical results is to understand the dynamics of load accumulation during range expansion when selfing evolves. A potential major benefit of selfing is the opportunity for purging due to increased homozygosity. Theory predicts that increased homozygosity should lead to efficient removal or reduction of lethal mutations (Kirkpatrick and Jarne 2000; Hedrick 2002), but our simulation results show that expansion load always accumulates at similar levels at range edges, regardless of selfing rate and with equivalent severity to outcrossers. This seems to suggest that purging due to selfing offers no additional benefit during species range expansion. However, when looking at the distribution of effect sizes for variants segregating within populations, we detect significant effects of purging unique to selfers whereby lethal-effect alleles are successfully and rapidly removed from the population. Purging was most pronounced in obligate selfers, where within only 30-150 generations lethal alleles are removed from the population and remain at low levels for the remainder of the simulations (Figure 2B). Examining the distribution of mutational effect sizes at the end of the simulations also shows that selfers exhibited major reductions in lethal alleles (Figure 3), to a much greater degree than the reduction of lethals obtained by outcrossers.

Purging does not, however, allow these populations to escape the burden inflicted by expansion load and their demographic past. Load still accumulates in all population expansions regardless of mating system, but how this load is expressed in terms of number and effect size of variants differs among mating systems. The overall burden experienced by selfing populations consists of more small-effect deleterious variants, which accumulate to a greater extent as compared to obligate outcrossing populations. Previous simulation studies have also highlighted how small effect variants are much more difficult to purge (Wang *et al*. 1999; Willis 1999), and therefore can still result in an accumulation of expansion load. How this genetic architecture underlying expressed load differs may have important impacts on how selection and recombination interact as populations adapt in the future. This burden represented by many small effect loci should also indicate that inbreeding depression is minimized or removed if populations continue to self, as previously described by (Pujol *et al*. 2009).

In our empirical *A. alpina* results we found similar signatures of both load accumulation and genetic purging in expanded populations. The recessive load model indicates that French and Swiss expanded populations have accumulated genetic load, through higher counts of putatively deleterious sites. Moreover, the additive model shows a small decrease in deleterious counts for our expanded populations. This suggests that negative selection has purged some diversity from these populations, since otherwise allele counts should remain at constant levels across all populations if only genetic drift is acting and not selection (Peischl and Excoffier 2015). However, our empirical DFE inferences only detected minor trends of reduced proportions of sites in the most deleterious class for expanded Swiss and French populations and equivalently support a bimodal-shaped DFE reported in Laenen *et al*. (2018). Whether this reflects true minor differences in the DFEs among these populations, or a lack of proper inferential ability is difficult to know. Comparing genetic load in populations with complex population demographic histories is still challenging (Simons and Sella 2016; Brandvain and Wright 2016), largely because dominance coefficients are unknown and different DFEs may respond differently to estimation accuracy (Gilbert *et al*. 2022) or simply differently to demographic change (Glémin 2003; Balick *et al*. 2015). Other approaches which attempt to directly infer mutational effects require intensive and in-depth analyses, emphasizing the need for improved inference methods or further in-depth investigation to accurately and efficiently identify effect sizes of variants in natural populations and therefore gain a complete picture of the genetic architecture of mutation load. Still, our 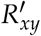 analyses provide additional support for our load estimates across populations, showing an accumulation of derived deleterious alleles relative to neutral alleles in all expanded populations relative to Italian populations. Only further along our expansion axis, in Swiss populations, is there some evidence that further accumulation of load is purged due to selfing.

The concurrence between our simulated and empirical results gives striking insights into the interactions of demographic change due to range expansion with recombination and diversity changes due to self-fertilization. We observe that during the process of range expansion, newly colonized demes are highly subject to drift and founder events, as extensively discussed in previous studies on range expansions (Excoffier *et al*. 2009; Slatkin and Excoffier 2012; Peischl *et al*. 2013, 2015). Furthermore, the added impact of self-fertilization exacerbates losses of heterozygosity at expanding range edges. This beneficially allows for the removal of recessive lethal alleles, but such large losses of diversity should otherwise hinder future adaptation. Though our simulations indicate that this loss of diversity can be recovered after the expansion front has passed, when migration and population growth allow for increased efficiency of selection in larger and more diverse populations, as previously described in Gilbert *et al*. (2018). A novel insight from our results is that this recovery is much slower for selfing populations, supporting the widely-held idea that selfing should only be favored at range edges and that outcrossing may replace selfing after a range expansion has occurred. Given the result that our empirical populations still exhibit signatures of genetic load is then equally interesting, since these populations are expected to have had many generations for recovery since they were directly on an expanding edge. The dynamics of when selfing evolves and can be favored across a species range is examined *in silico* by Encinas-Viso *et al*. (2020), showing that unless recombination rates are high enough, outcrossing individuals will outcompete selfing populations once the expansion edge has passed. Whether our sampled alpine populations populations have also began shifting back towards increased outcrossing is currently unknown and an avenue of investigation which will be interesting to pursue in the future.

Studying range expansions in plant species offers unique insights into the combination of mating system evolution combined with the evolutionary processes occurring during species range expansions. Previous work in *A. alpina* has also evidenced increased load with high selfing and bottlenecks in Scandinavian populations (Laenen *et al*. 2018), however we highlight previously unidentified evidence for purging of strongly deleterious alleles in intermediate to highly selfing continental populations within the French and Swiss Alps, in addition to expansion load still incurred. Our results are also similar to those found in other plant range expansions where selfing is observed at the range edge. Notably, this is the case in *A. alpina*’s close relative *Arabidopsis lyrata* (Willi *et al*. 2018) as well as in *Mercurialis annua* (González-Martínez *et al*. 2017) where expansion load has been indicated. Our study has uniquely also identified signature of purging due to selfing, which is a known expectation from theoretical predictions (Glémin 2003, 2007), but to our knowledge not thoroughly investigated in empirical systems. Future studies could likewise benefit from direct estimates of fitness across the species range, through crosses and common garden studies.

Population genetic simulations help us to better understand interactions of effects that are difficult to assess or disentangle in empirical populations. Here, we have only explored a finite parameter space and constrained our simulations to simplified demographic models. Since we were only interested in the eventual signatures resulting from selfing evolution during a range expansion, we modeled the loss of self-incompatibility as a sudden shift in the probability of selfing at one location on the landscape. However, in nature, the shift to self-fertilization is expected to occur gradually over time, e.g, due to a reduction in S-allele diversity (Charlesworth and Charlesworth 1979; Vallejo-Marín and Uyenoyama 2004; Porcher and Lande 2005; Encinas-Viso *et al*. 2020). Even with our sudden evolution of selfing imposed in the middle of the landscape, we expect that the same observed qualitative results of purging strong-effect recessive deleterious alleles and loss of heterozygosity would still occur, just more gradually through time. The intermediate rate of selfing we tested could also be considered an earlier transitional state of a species range expansion on its way to evolving higher selfing rates. In a gradual shift to selfing, initial S-allele diversity would be reduced but outcrossing still frequent, and intermediate selfing rates would be a transient state as populations shift to higher selfing and faster expansion. While we focused on the speed and purging benefits of selfing during a range expansion, we did not address a potential third factor impacting expansions: the necessity to locally adapt to unfamiliar environments. Populations must often adapt to novel or fluctuating environments during expansion, e.g., during glacial cycles (Hewitt 2004) or as soil conditions change over altitude or photoperiod conditions change over latitude. Adaptation requires sufficient genetic variation to match the local environment sufficiently for population growth to be sustainable (Kirkpatrick and Barton 1997; Polechová and Barton 2015). For populations that expand to follow an environment they are already adapted to, this difficulty is less relevant. For example, species expanding post-glaciation are believed to have followed the receding ice sheets as suitable habitat that they were pre-adapted to was slowly revealed. However, it is still likely that some aspect of environmental conditions are always novel as organisms move over space, necessitating some level of adaptation. Our results importantly highlight how the DFE is expected to differ among outcrossed versus selfed expanding populations, creating contrasting genetic architectures within the genome. Such differences in genetic architectures, i.e., few large-effect or many small-effect loci, for adaptive and maladaptive sites along with differing effective recombination rates across selfing rates are likely to interact with adaptation over changing landscapes and result in different adaptive potentials among populations. And in the future, as anthropogenically-induced climate change causes more rapid changes across the landscape, the likelihood of being able to track moving environmental optima is expected to become more difficult, necessitating more rapid adaptation and emphasizing the importance of studying range expansions and shifts and the evolutionary processes involved.

## Conclusions

Range expansions are known to increase genetic drift and fixation of deleterious alleles, reducing fitness as a consequence. Self-fertilization further reduces *N*_*e*_, which allows for a higher rate of fixation of weaker deleterious mutations compared to outcrossers. However, as predicted by Glémin (2007) this process can also allow for short-term purging. We investigated whether this purging is realized during species range expansions and if selfing can thus be beneficial in this evolutionary context. We described two significant factors in our simulations: first, the purging of lethal alleles is indeed observed in selfing populations, and second, this purging is not sufficient to prevent the fitness loss incurred by expansion load. Weak effect mutations accumulate to a larger extent due to the range expansion, leaving a visible signature in the DFE. Furthermore, in natural populations of *A. alpina*, we see consistent effects of purging as well as load accumulation despite the evolution of selfing. Together, this demonstrates that self-fertilization can alter the signature of genetic load in expanded populations, and identifies purging as an additional benefit of selfing along with reproductive assurance. Future studies in empirical systems will hopefully be able to distinguish expanded outcrossing versus expanded selfing populations to further validate our results, as much remains to be learned of the interaction between mating system evolution and demographic history of populations. Improved understanding of these important processes will be vital for further insight into how natural populations will (or will not) be able to disperse and adapt in the face of global climate change and anthropogenic forces experienced in natural habitats.

## Material and Methods

We conduct simulations of a species range expansion and compare to an empirical dataset from the plant *Arabis alpina* to understand the dynamics of purging and mutation load accumulation in a system where self-fertilization has evolved. To understand whether selfing acts as an evolutionary advantage during expansion by purging deleterious alleles that otherwise accumulate, we focus on tracking genetic load in both simulated and empirical data. Though simplified from reality, our simulations have the important advantage of knowing true fitness and mutational effects within every individual to best understand the dynamics of load accumulation and purging during range expansion.

### Simulations

To simulate a range expansion with a shift in mating system we conducted individual-based, forward time simulations, using a non-Wright-Fisher model in SLiM v3.7.1 (Haller and Messer 2019). We modeled the range expansion across a one-dimensional, linear landscape of 50 demes with a stepping-stone migration model (Figure 1A). Each simulation started with a single initial core deme populated with individuals that then underwent repeated bottlenecks and founder events as they colonized the remaining empty 49 demes. The core population was initiated at carrying capacity *K* = 5000, and prior to expanding we ran a burn-in for 4*N* generations. Generations were discrete and non-overlapping, and after the burn-in was complete we opened the landscape for expansion, introducing migration that allowed individuals to move into either adjacent deme. We defined a forward migration rate of *m* = 0.05 per generation and reflecting boundaries at the ends of the landscape in the core and deme 50. All subsequent demes outside the core had a carrying capacity of *K* = 200. Once the last deme reached carrying capacity and 100 additional generations passed, we stopped the simulation.

To test the effect of increased self-fertilization during the expansion we conducted a set of obligately outcrossing simulations to serve as a null model for range expansion without the additional impact of uniparental inbreeding arising from selfing. We then compared to three different simulated scenarios where selfing begins halfway through the expansion, in deme 25. In demes 25 − 50 of these selfing simulations, we set the self-fertilization rate *σ* to either 0.5, 0.95, or 1. We replicated every parameter combination 20 times for a total of 80 simulations across all three selfing rates and the obligate outcrossers. In a given deme, individuals to be selfed were chosen with probability *σ* each generation. We modeled logistic population growth with a Beverton-Holt model, where the expected number of total offspring per deme for the next generation is given by 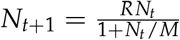, where 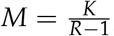, growth rate *R* = 1.2, *N*_*t*_ is the deme’s census size in the current generation *t*, and *K* is the carrying capacity of the focal deme. For each parent, we expected less fit individuals to produce fewer offspring and thus implemented a fecundity selection model, where the expected number of offspring for individual *i* is approximately Poisson distributed (Peischl *et al*. 2015).

Each individual was modeled as a diploid genome consisting of 1× 10^7^ base pairs (bp) with a recombination rate of 1×10^−8^ per bp per generation. We simulated neutral, beneficial, deleterious, and lethal mutations at a per base pair mutation rate of 7 × 10^−8^ per generation occurring at relative proportions of 0.25, 0.001, 0.649 and 0.1, respectively. For deleterious and beneficial mutations, selection coefficients were drawn from an exponential function with mean -0.001 or 0.01, respectively, and lethal alleles had a selection coefficient of -1. Dominance coefficients were set to *h* = 0.3 for beneficial and deleterious alleles, and 0.02 for lethal mutations. Individual fitness in SLiM is calculated multiplicatively across all mutations an individual possesses, as drawn from these distributions for effect size and dominance coefficient. In a supplementary set of simulations we tested for the effect of full additivity using *h* = 0.5 for non-lethal mutations. These simulated parameters for the distribution of selection and dominance coefficients reflect partial dominance of deleterious alleles and more recessive lethal alleles, as described in the literature for the current best knowledge of mutational distributions in nature (Keightley 1994; Eyre-Walker and Keightley 2007; Halligan and Keightley 2009; Agrawal and Whitlock 2011).

We recorded fitness and calculated summary statistics during the expansion to track the impact of demographic change in combination with selfing rates. In every deme we measured nucleotide diversity for neutral variants, *π*, mean observed heterozygosity along the genome, *H*, counts of lethal and deleterious alleles and recorded mean fitness, 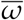, every five generations. This allowed us to compare changes in fitness and allele counts over time, contrasting them with the same statistic 100 generation in the past. We also examined changes in these summary statistics in specific locations across the landscape during and after the expansion had completed: the core (deme 1), the deme prior to the mating shift (deme 24), the deme ten demes past the facultative mating shift (deme 35, to avoid effects of migration from outcrossers), and the end of the landscape (deme 50). We characterized the composition of load in core and edge populations after the expansion by examining the realized distribution of selection coefficients. To do this, we categorized selection coefficients in four discrete bins (*s* ∈ { (0.0001, 0), (0.001, 0.0001], (1, 0.001], −1) }). To further characterize load, we calculated the proportion of fixed deleterious alleles, and applied models often used to compare approximated genetic load in empirical populations (Simons and Sella 2016) to our simulated data: we estimated additive load by counting the total number of deleterious alleles per individual, assuming *h* = 0.5, and recessive load by counting the total number of homozygous deleterious loci per individual, assuming *h* = 0. We then compared these values with realized fitness, all of which are known for the simulations.

### *Arabis alpina* dataset

We compared our theoretical results to an empirical dataset of *Arabis alpina* by combining publicly available data (Laenen *et al*. 2018; Rogivue *et al*. 2019) with newly sampled and sequenced genomes. Our dataset focused on sampling four regions with four populations each, consisting of 15-18 individuals. One exception is northern Italy where two nearby populations (Ca & Gf) of 8 individuals each contributed to five total populations from the region. Sampling spans the range expansion from southern Italy north into the French and Swiss Alps and capturing the transition in mating system from outcrossing to selfing. We collected leaf tissue on silica gel from 198 wild *A. alpina* plants in the Apennine Mountains in central Italy, the Apuan Alps in northern Italy, and the western Alps in France during the summer of June 2021. We extracted DNA with the Qiagen DNeasy Plant Mini Kit (Qiagen, Inc., Valencia, CA, USA) and constructed libraries using Illumina TruSeq DNA PCR-Free (Illumina, San Diego, CA, USA) or Illumina DNA Prep, and sequenced on a Illumina NovaSeq 6000 (paired-end). All sampled individuals are described in more detail in Supplementary Table 1 and are available publicly at NCBI SRA accession PRJNA773763. We combined this dataset with previously published *A. alpina* short-read genomes of 306 individuals sampled from Switzerland (Rogivue *et al*. 2019) and 36 sampled widely across Europe (Laenen *et al*. 2018). For quality control of the reads, we used FastQC (http://www.bioinformatics.babraham.ac.uk/projects/fastqc) and MultiQC (Ewels *et al*. 2016). We trimmed reads using trimmomatic 0.39 (Bolger *et al*. 2014) and aligned them to the *A. alpina* reference genome (Jiao *et al*. (2017), version 5.1, http://www.arabis-alpina.org/refseq.html) using `bwa mem` 0.7.17 (Li and Durbin 2009). We calculated coverage for the whole dataset with mosdepth (Pedersen and Quinlan 2018), averaging at 13.98 (18.61 for new samples, Supplementary Table 1). We called variant and invariant sites using freebayes 1.3.2 (Garrison and Marth 2012). Additional filters were applied in bcftools (Danecek *et al*. 2021), retaining only sites with a maximum missing fraction of 0.2, and removing any variant sites with estimated probability of not being polymorphic less than phred 20 (QUAL>=20). Finally, we removed 13 individuals with greater than 30% missing calls or low coverage (Br22, Br06, Cc05, St15, Am01, Br18, Br24, Ma28, Pa9, Pi9, Pi95, Pi40, Ma97). The final dataset combined had 3,179,432 SNPs, with 43,268,666 invariant sites for 527 individuals from 31 populations, which includes 191 individuals of the 17 newly sampled populations in the Italy-Alps expansion zone.

### Population genetic analyses

We inferred the ancestral state of alleles using the close relative of *A. alpina, Arabis montbretiana*, by aligning the reference sequences of *A. alpina* with *A. montbretiana* (Madrid *et al*. 2021) `using last` (Kiełbasa *et al*. 2011). To confirm that our samples from across Europe matched the expected population structuring based on known demographic history, we ran `admixture` v1.3.0 (Alexander *et al*. 2009) from *K* = 2 to *K* = 15 on the full sample set but with SNPs pruned for LD using `bcftools +prune` (Danecek *et al*. 2021, *R*^2^ cutoff 0.3 in a window of 1000 sites). We calculated nucleotide diversity per population in 1Mbp windows using pixy (Korunes and Samuk 2021, version 1.2.6.beta1, 10.5281/zenodo.6032358) and inbreeding coefficients for each individual with ngsF (6 iterations, Vieira *et al*. 2013). To format the input file for ngsF, we randomly sampled 100,000 biallelic SNPs and extracted genotype likelihoods using bcftools (Danecek *et al*. 2021).

To calculate putative genetic load, we first predicted derived, deleterious alleles with SNPeff (Cingolani *et al*. 2012). SNPeff estimates how deleterious a variant may be based on whether its mutation causes an amino acid change, at varying levels of importance (Cingolani *et al*. 2012). We used the categories “nonsense” and “missense” as the definition for deleterious mutations from SNPeff, “none” and “silent” annotations were used as neutral predictions and “LOF” annotations as loss-of-function mutations (“LoF”) after running the program with the `-formatEff` option. Using these annotations, we calculated 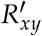 as described in Do *et al*. (2015) with *R*_*xy*_ for derived allele counts of LoF or deleterious sites over 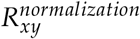 for neutral sites to avoid reference bias. Further, we estimated jackknife confidence intervals using pseudo values from 100 contiguous blocks and assuming normal distributed values. We also estimated genetic load from SNPeff predictions, using the same recessive and additive models as in the simulations, by counting either the total number of deleterious loci or derived alleles at deleterious sites. Finally, we estimated the empirical DFE for every population using `fitdadi` (Kim *et al*. 2017) in 100 replicated runs (see Figures S5, S8 for supplementary methods and results).

## Data and code availability

Statistical analyses were conducted in R v.4.1.3 (R Core Team 2018), unless otherwise specified. Genetic data is archived at NCBI SRA (accession PRJNA773763). Code and simulation output is available on GitHub (https://github.com/LZeitler/selfing_expansion).

## Acknowledgments

We thank Drs. Marco Andrello, Michele Di Musciano, Marta Binaghi, Paola Morini, Marco Caccianiga, Alessandro Alessandrini, Rodolfo Gentili, and Enzo Bona for essential help in finding wild populations of *A. alpina* to sample in Italy and France. We thank Dr. Pamela Nicholson and the team at the Bern NGS Platform for sequencing and troubleshooting help as well as Ryan Gutenkunst for help troubleshooting the dadi analyses. We thank Stephan Peischl for useful feedback on the manuscript and Xuejing Wang for statistical advice. This research was funded by Swiss National Science Foundation Ambizione grant #PZ00P3_185952 to KJG.

## Author Contributions

LZ contributed to further developing the study’s idea, designed, wrote, and ran the simulations, led the empirical analyses, participated in sample collection and DNA extractions, and wrote the manuscript. CP contributed to further developing the study’s idea and to editing the manuscript. KJG conceived the original idea for the study, coordinated the study, contributed to designing the simulations, participated in sample collection and DNA extractions, and contributed to writing and editing the manuscript.

## Supplementary

**Figure S1:**
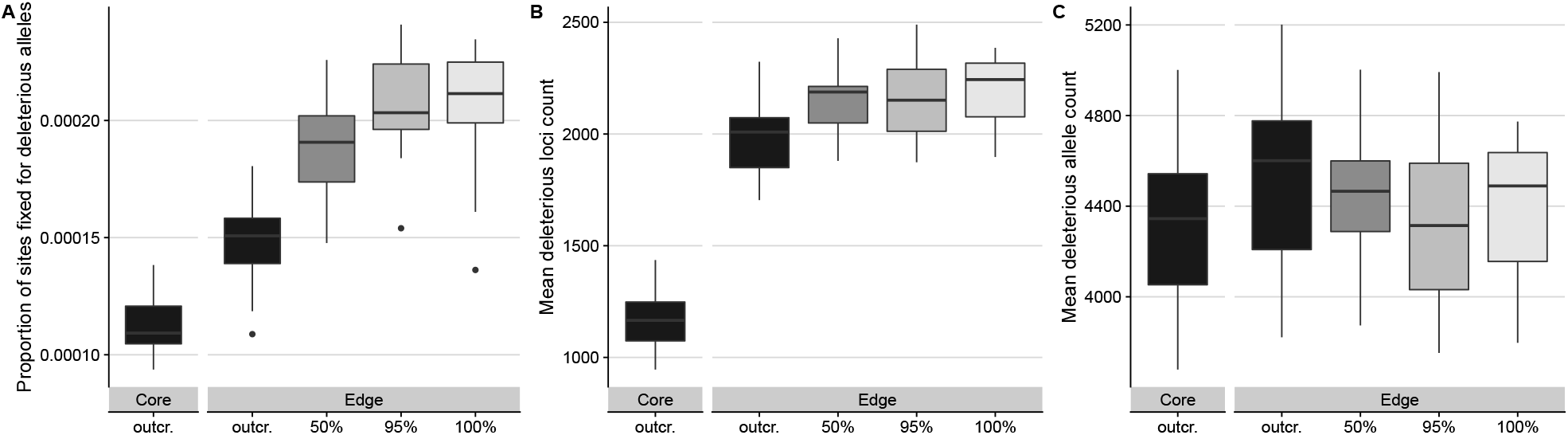
The proportion of sites fixed for deleterious alleles (A), the mean counts of deleterious loci (B), and the mean counts of deleterious alleles (C), all assessed at the end of the simulations for core and edge populations across selfing rates. Whiskers indicate the 1.5 interquartile range.

**Figure S2:**
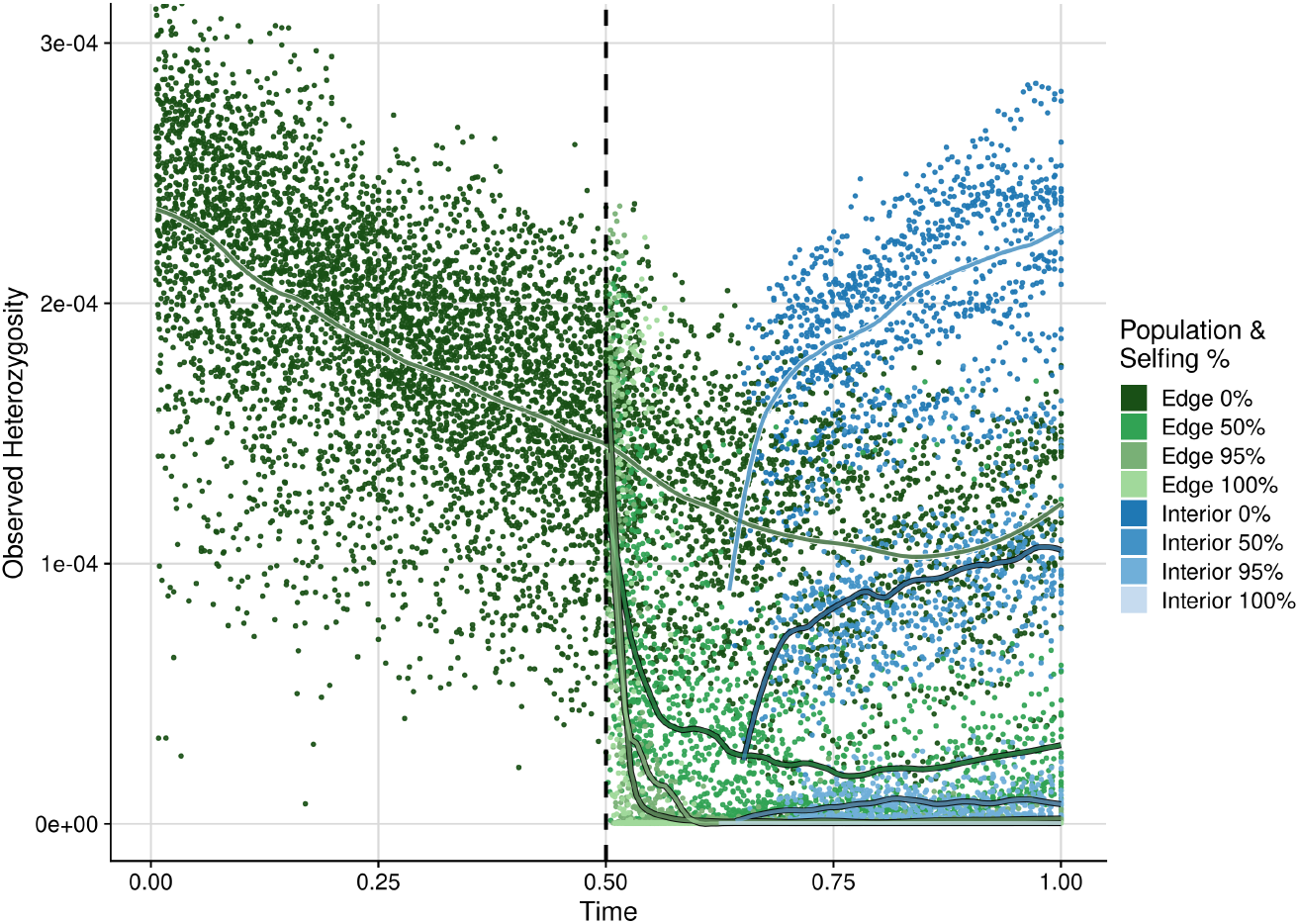
Trajectories for the mean observed heterozygosity over relative time, as described in Figure 2A, but now including all simulated selfing rates.

**Figure S3:**
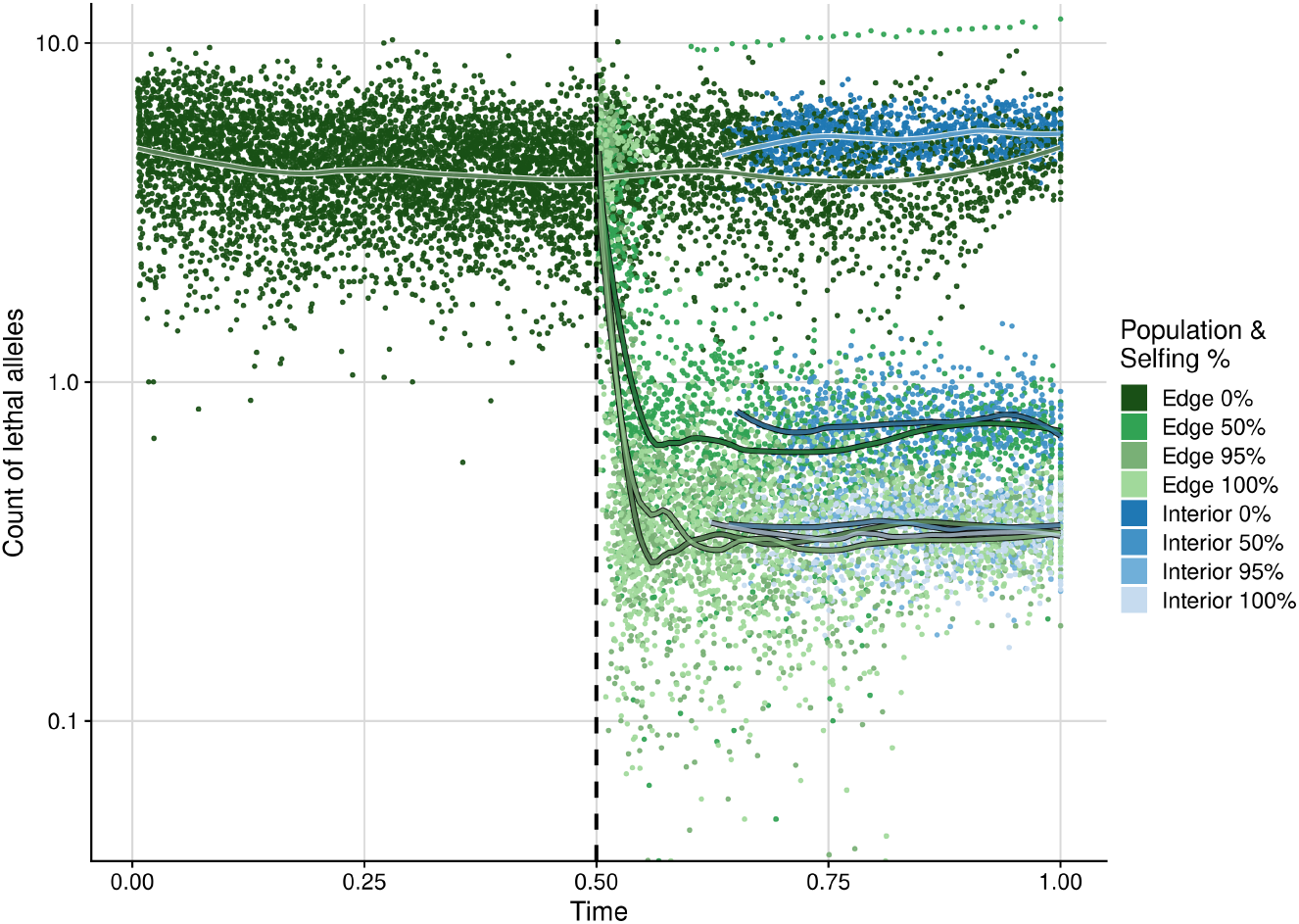
Trajectories for the mean count of lethal alleles over relative time, as described in Figure 2B, but now including all simulated selfing rates.

**Figure S4:**
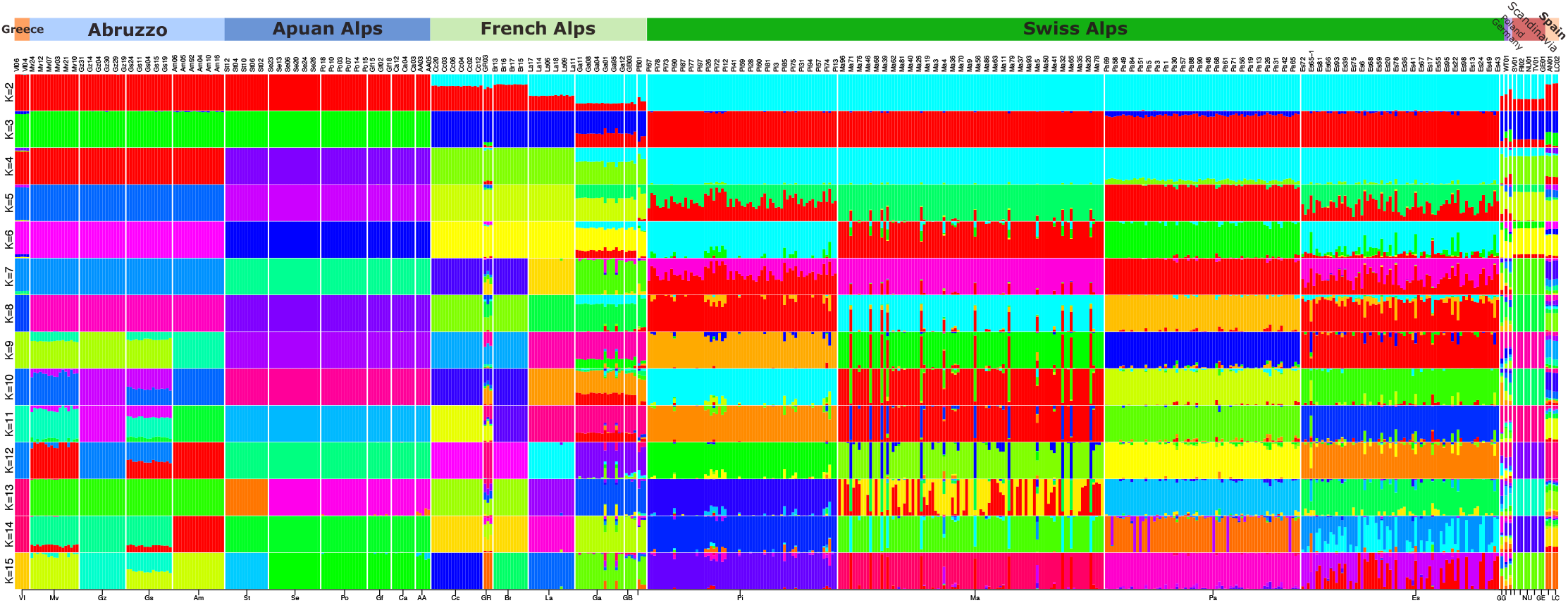
Results from *K* = 2 to *K* = 15 from admixture analyses run on the combined empirical dataset across Europe. The lowest CV error is for *K* = 14, however it is most useful to compare the populations structure across values of *K* to see how well this matches known geography and demographic history of the populations. We observe clean distinctions among our geographic regions sampled (indicated above the bar plots), with evidence for some gene flow across geographic space as one observes higher *K* values.

**Figure S5:**
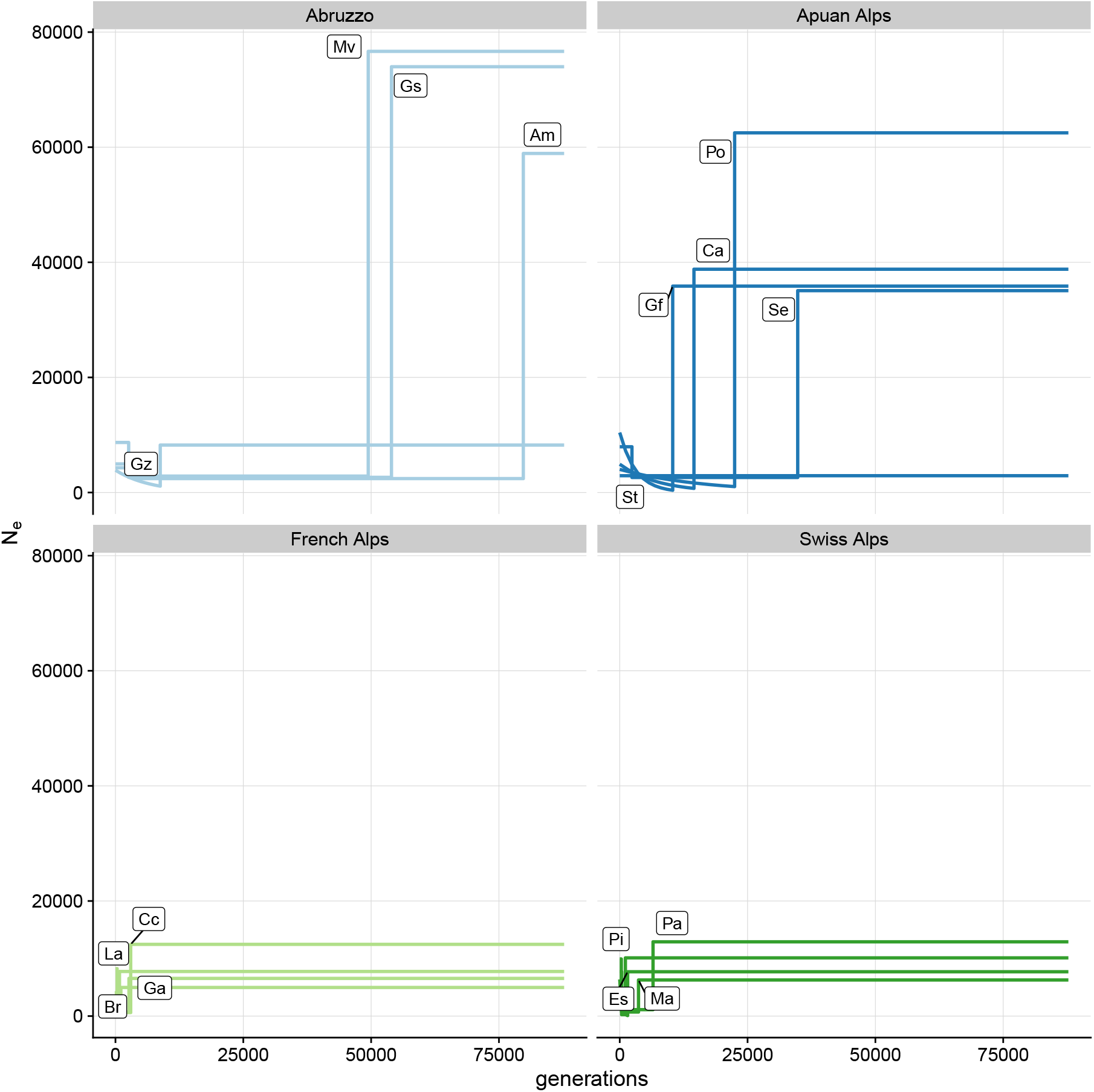
We inferred the demographic history of each of our newly sampled populations of *A. alpina* along with the densely sampled Swiss populations using dadi. This is a necessary step to account for the demography when inferring the DFE with fitdadi. This also allowed us to confirm if this newly inferred demographic history is consistent with past studies in *A. alpina*. The best-fitting models for our populations, based on AIC, were “bottlegrowth” models, indicating a past bottleneck followed by exponential growth (Es, Ca, Gf, Gz, Po), three epoch models, indicating a bottleneck followed by a sudden size change (Pa, Pi, Ma, Am, Br, Cc, Ga, Gs, La, Mv, Se), and the standard neutral model (St). Populations St and Gz were the only instances where competing models fitted approximately equally well (see Supplementary Table 2), therefore results for these population should be interpreted with caution. With the exception of Es, all Alpine populations best fit to three epoch models. Central Italian populations (light blue) show the most historic bottlenecks and the largest ancestral populations sizes. This is consistent with this region of highly outcrossing plants being subject to the last glacial maximum. Northern Italian populations (dark blue) show more recent bottlenecks and reduced ancestral sizes relative to central Italy, potentially reflecting their expansion northward. French and Swiss Alpine populations both showed the most recent bottlenecks and the smallest historic population sizes, consistent with both their shift to selfing and their more recent range expansion. Depleted genetic diversity along the axis of an expanding species range is expected (Pujol *et al*. 2009; Peischl *et al*. 2013, 2015), as is decreased *N*_*e*_ due to inbreeding and thus loss of diversity (Keller and Waller 2002; Charlesworth and Willis 2009). These demographic inferences thus match our understanding of both the mating system shift and the range expansion that these populations experienced.

**Figure S6:**
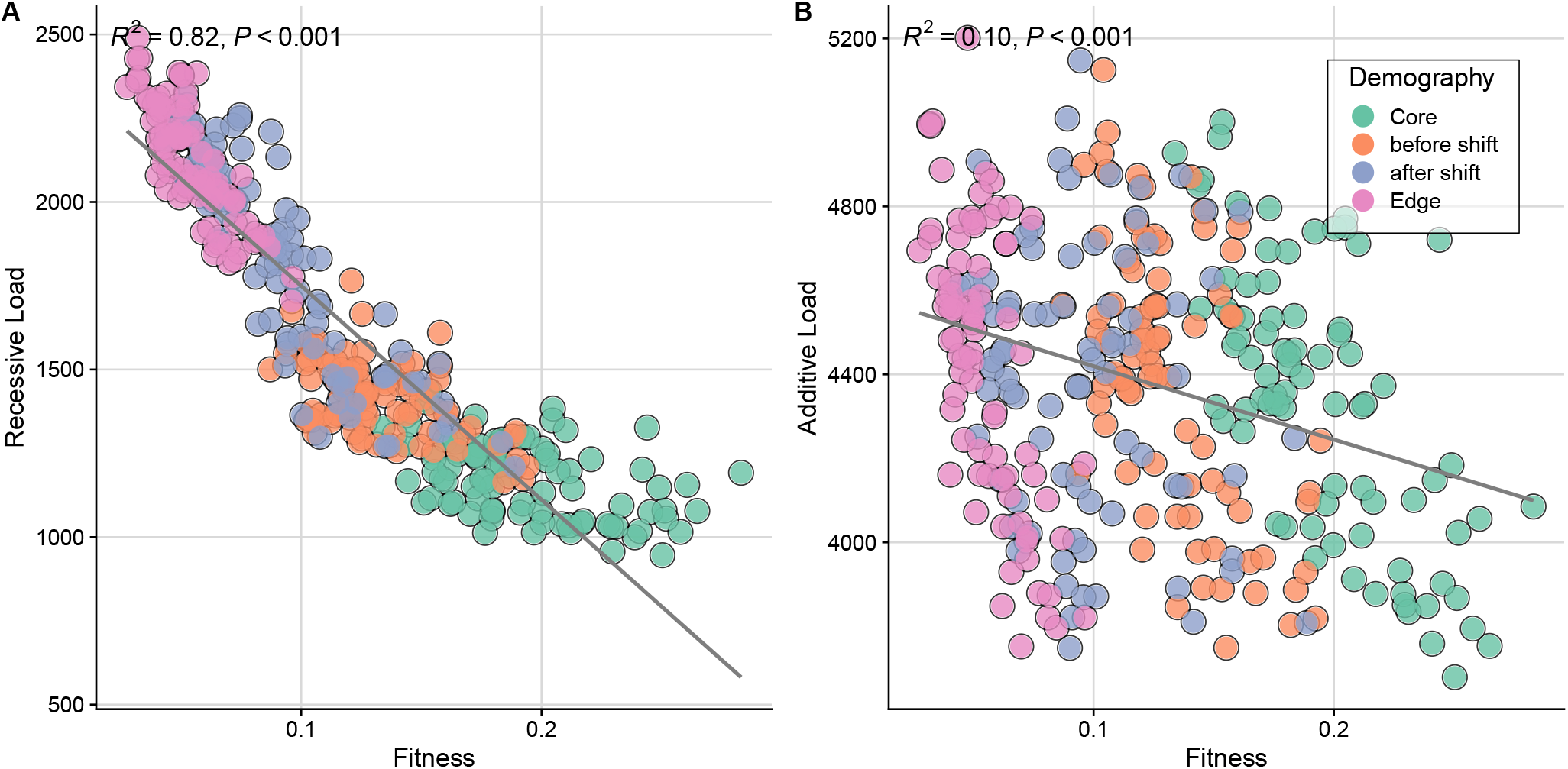
Observed (known) mean fitness from simulations for core (green), interior (orange, purple) and edge (pink) demes compared to the inverse of the count of deleterious loci (A), both after the range expansion is complete. The count of deleterious loci serves as a model for recessive load, which we find best correlates to fitness, compared to the additive model (B), where load is predicted by counting alleles. Results are for simulations with *h* = 0.3 for non-lethal deleterious mutations.

**Figure S7:**
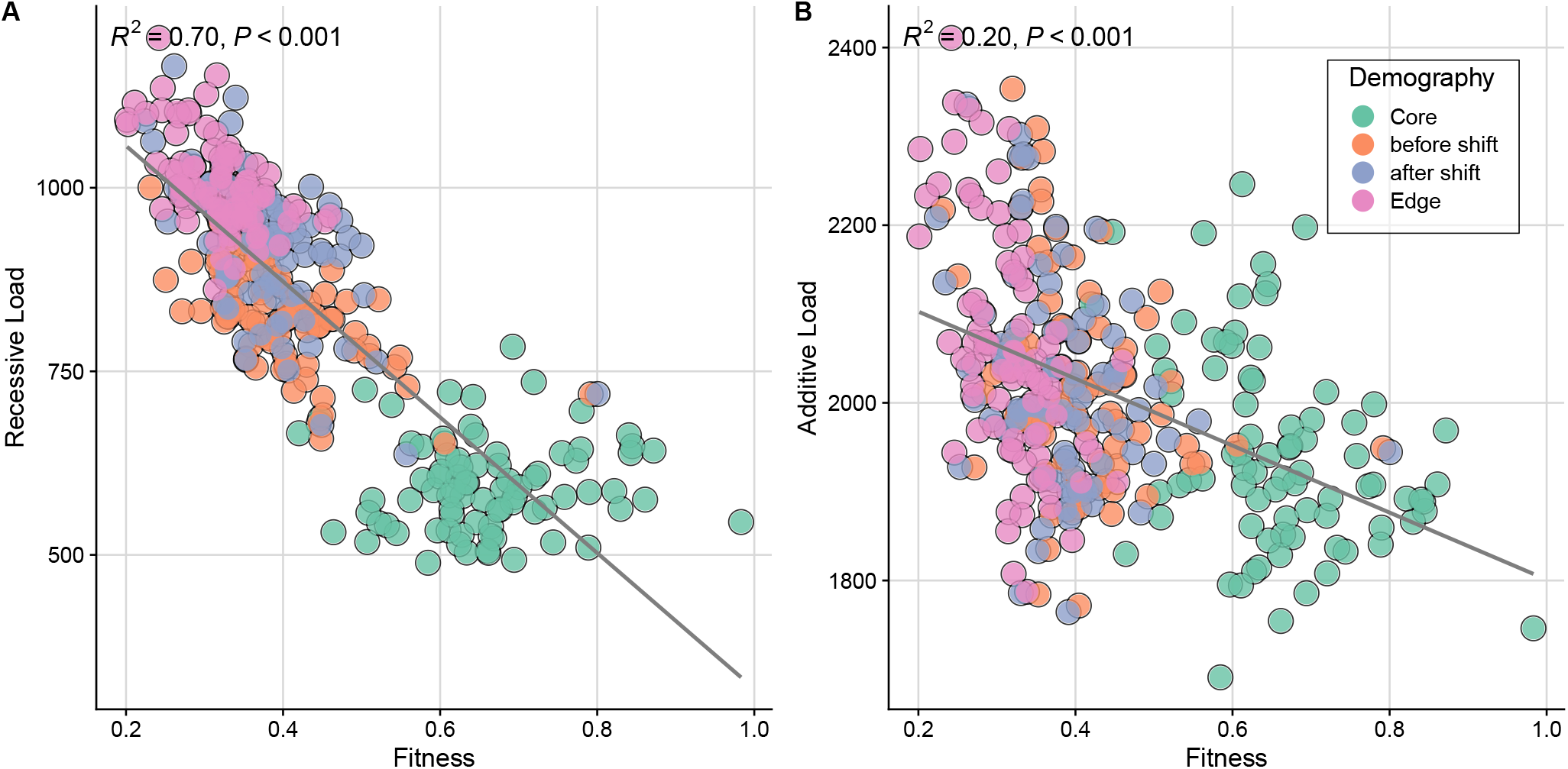
Recessive (A) and additive (B) genetic load compared with known simulated fitness to infer load when all non-lethal deleterious mutations are perfectly additive (*h* = 0.5). Data is from a supplementary set of simulations with these dominance parameters. This repeats the same analyses as Figure S6, except now for simulations with additive mutations. This result again finds that the recessive model predicts load better (*R*^2^ = 0.70, *P* < 0.001) than the additive model (*R*^2^ = 0.20, *P* < 0.001).

**Figure S8:**
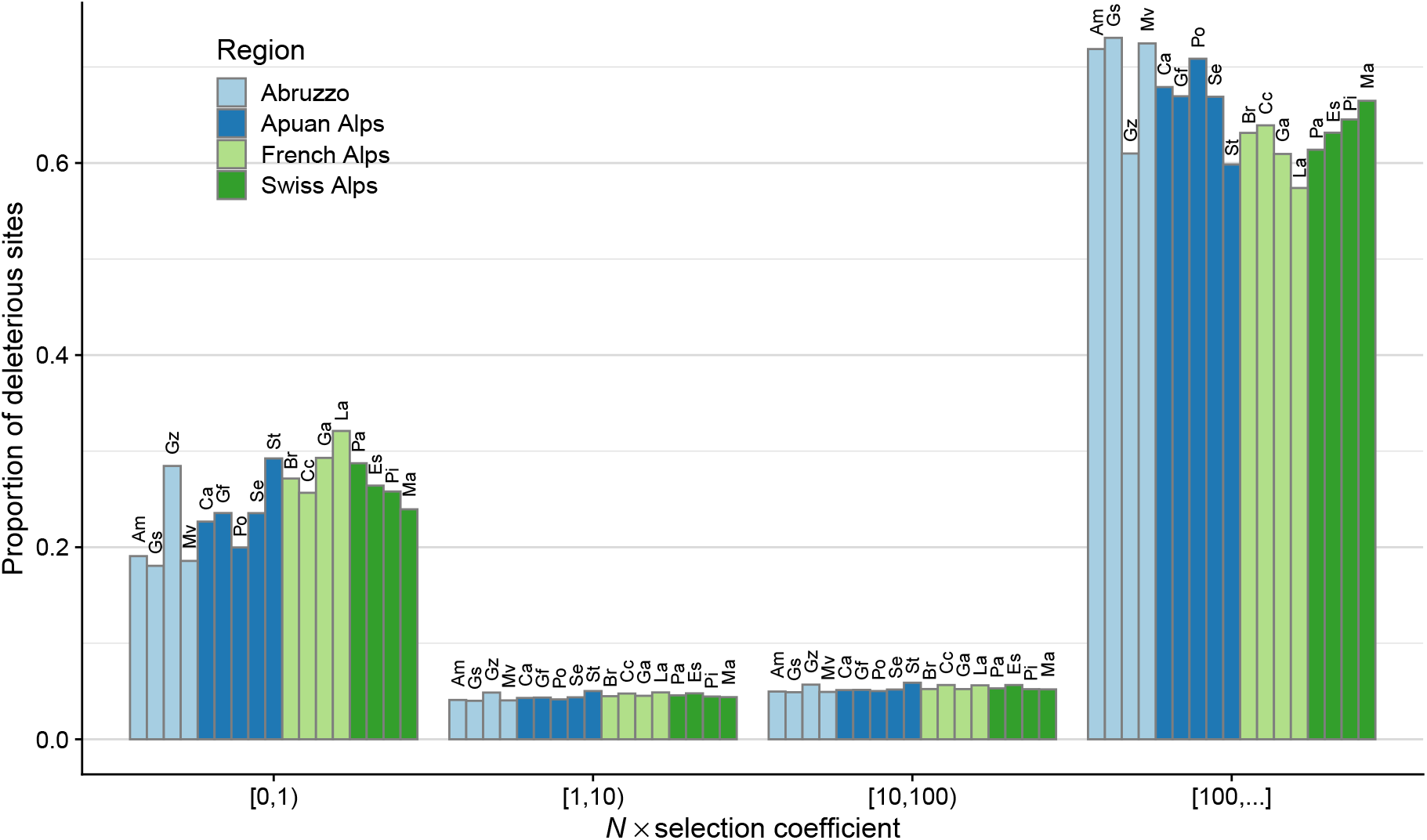
We inferred the DFE of each *A. alpina* population in the Italy-Alps expansion zone using fitdadi (Kim *et al*. 2017) from dadi (Gutenkunst *et al*. (2009) in python 3.8.12. We used the SNPeff annotation to construct polarized site-frequency spectra for neutral and deleterious sites after subsampling to a maximum population size of 20 individuals. To estimate demographic parameters, we tested the default single population demographic models (standard neutral model, two-epoch, growth, bottlegrowth, threeepoch) and two models accounting for inbreeding (standard neutral with inbreeding, two-epoch with inbreeding). We assumed a per base pair mutation rate of *µ* = 7 × 10^−9^ per generation, ran the default optimization for 100 replicates, and selected the best fit parameters within each demographic model based on likelihood and the best fit demographic model based on AIC. For fitdadi, we additionally assumed *L*_*ns*_/*L*_*s*_ = 2.85, dominance coeffient *h* = 0.3 and estimated the DFE for each model in 100 optimizations. We then chose the best-fit DFE optimization based on likelihood for each population for the previously chosen demographic model. DFE results from *A. alpina* populations across the Italian-Alpine range expansion for outcrossing populations from Abruzzo (light blue) and the Apuan Alps (dark blue) are compared to the selfing populations that have undergone range expansions into the French Alps (light green) and the Swiss Alps (dark green). We found mean proportions across all populations of 65.4% and 24.8% in the weakest and strongest selection classes, respectively. Less than 5% of sites segregated in the two intermediate selection classes. These proportions varied only marginally between core Italian populations (mean proportions 22.6% and 67.9% for weakest and strongest classes, respectively) and between expanded French and Swiss populations (means proportions 27.4% and 62.6% for weakest and strongest class).

**Figure S9:**
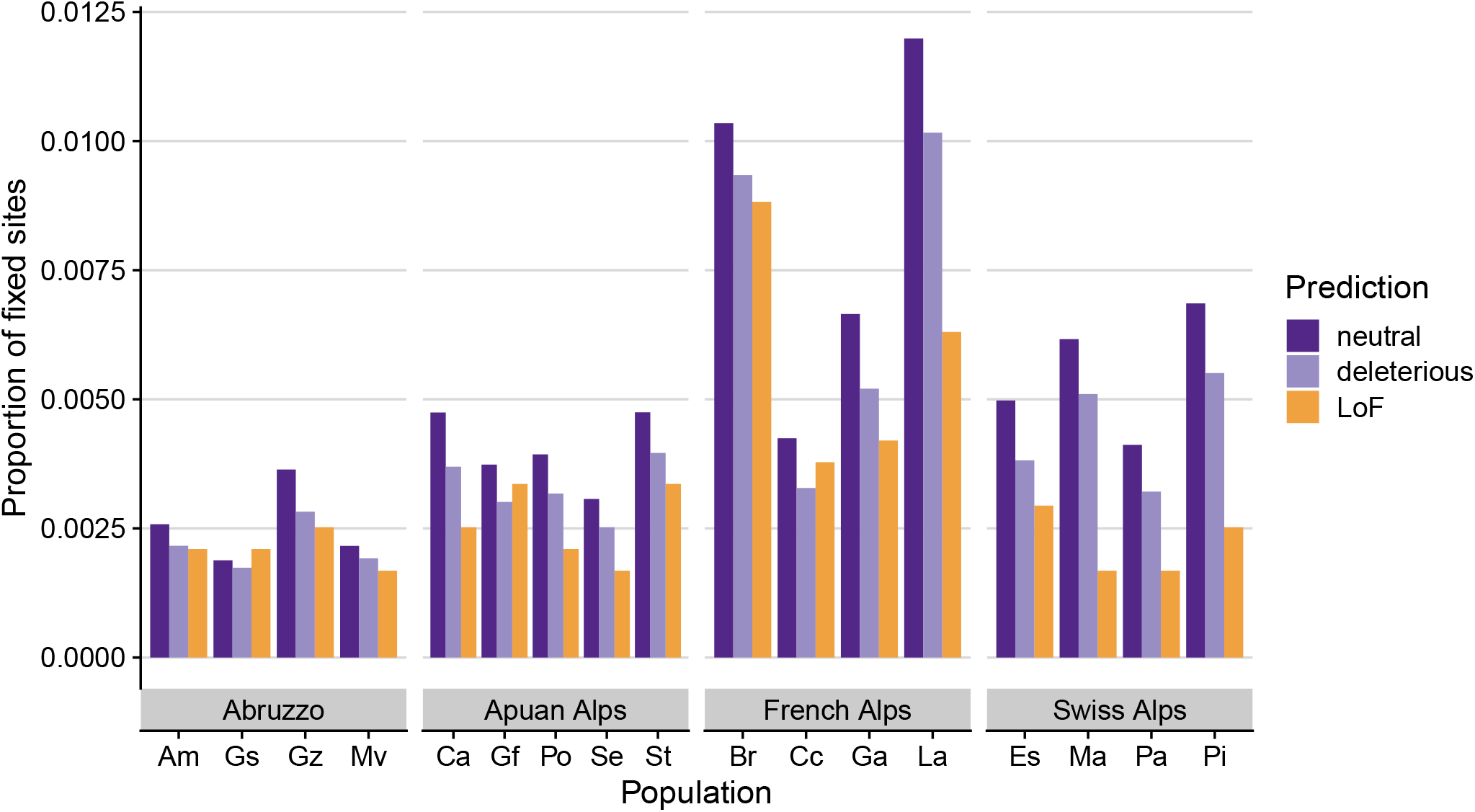
Fixation of predicted neutral (dark purple), deleterious (light purple) and loss of function (LoF, orange) sites per population. Y-axis shows the proportion of fixes sites in focal population and allele category. We found that neutral sites fixed at the highest proportions (mean 0.505%), while LoF sites were at the smallest proportions fixed (mean 0.314%), indicative of their highly deleterious effect. French populations Br and La had the highest overall fixation proportions of any class (0.948%), while samples from the Abruzzo region had the lowest (0.228%). Swiss population showed intermediate neutral fixation but LoF proportions similar to Italian populations.

